# The disordered JM motif in RTKs promotes classical DFG*_out_* conformation formation via dynamic effect

**DOI:** 10.1101/2025.10.28.684995

**Authors:** Xiaohui Chen, Hao Wang, Wenjian Li, Manjie Zhang, Bin Sun

## Abstract

Receptor tyrosine kinases (RTKs) are validated anti-cancer targets, and targeting their DFG*_out_* conformations represents a mainstream strategy for developing highly selective type II inhibitors. RTKs can adopt various DFG*_out_* conformations, but only the classical ones with a fully formed back pocket are structurally validated to accommodate type II inhibitors. However, such classical DFG*_out_* conformations are scarce, presenting a significant obstacle for selective RTK inhibitor design. Recently, a conserved disordered motif N-terminal to the kinase domain of RTKs, called the juxtamembrane (JM) motif, has been reported to regulate inhibitor binding to the DFG*_out_* conformation of VEGFR2, an RTK involved in angiogenesis. In this study, we performed extensive MD simulations to explore the impact of the disordered JM motif on the conformational space of the DFG motif in RTKs and its relationship to inhibitor binding at the active site of VEGFR2. We revealed that in VEGFR2, the disordered JM is highly dynamic and forms transient contacts with the kinase domain, and consequently fine-tunes the DFG*_out_* sub-conformational space to shift populations from non-classical to classical DFG*_out_* conformations. This dynamic model provides the structural basis underpinning the reported regulatory effects of JM on inhibitor binding toward VEGFR2. Addtionally, we demonstrated that in other RTKs beyond VEGFR2, the disordered JM similarly promotes classical DFG*_out_* conformations. Such role of JM is particularly favorable for creating druggable DFG*_out_* conformations that can be exploited for designing high-selectivity type II inhibitors.

## 1. Introduction

Receptor tyrosine kinases (RTKs) are a subfamily of the human kinome and are validated anti-cancer targets. They share a conserved domain organization, consisting of an extracellular ligand-binding domain, a transmembrane helix, and an intracellular kinase domain (**Fig. 1A**)^[1]^. Ligand binding to the extracellular domain induces receptor dimerization and autophosphorylation of cytoplasmic tyrosine residues, making RTKs key mediators of extracellular-to-intracellular signaling^[2]^. Abnormal RTK activation contributes to dysregulated angiogenesis and other oncogenic processes, validating RTKs as anti-cancer drug targets^[1]^. Small-molecule inhibitors targeting the kinase domains of RTKs represent a promising therapeutic strategy^[3]^. However, this approach is often limited by selectivity issues, which can lead to off-target side effects. This challenge arises because most FDA-approved RTK drugs are ATP-competitive inhibitors that bind the conserved kinase active site^[4]^, and the ATP- binding pockets across the more than 500 protein kinases in the human kinome are highly similar. Therefore, improving the selectivity of RTK inhibitors is critical.

**Figure 1.**
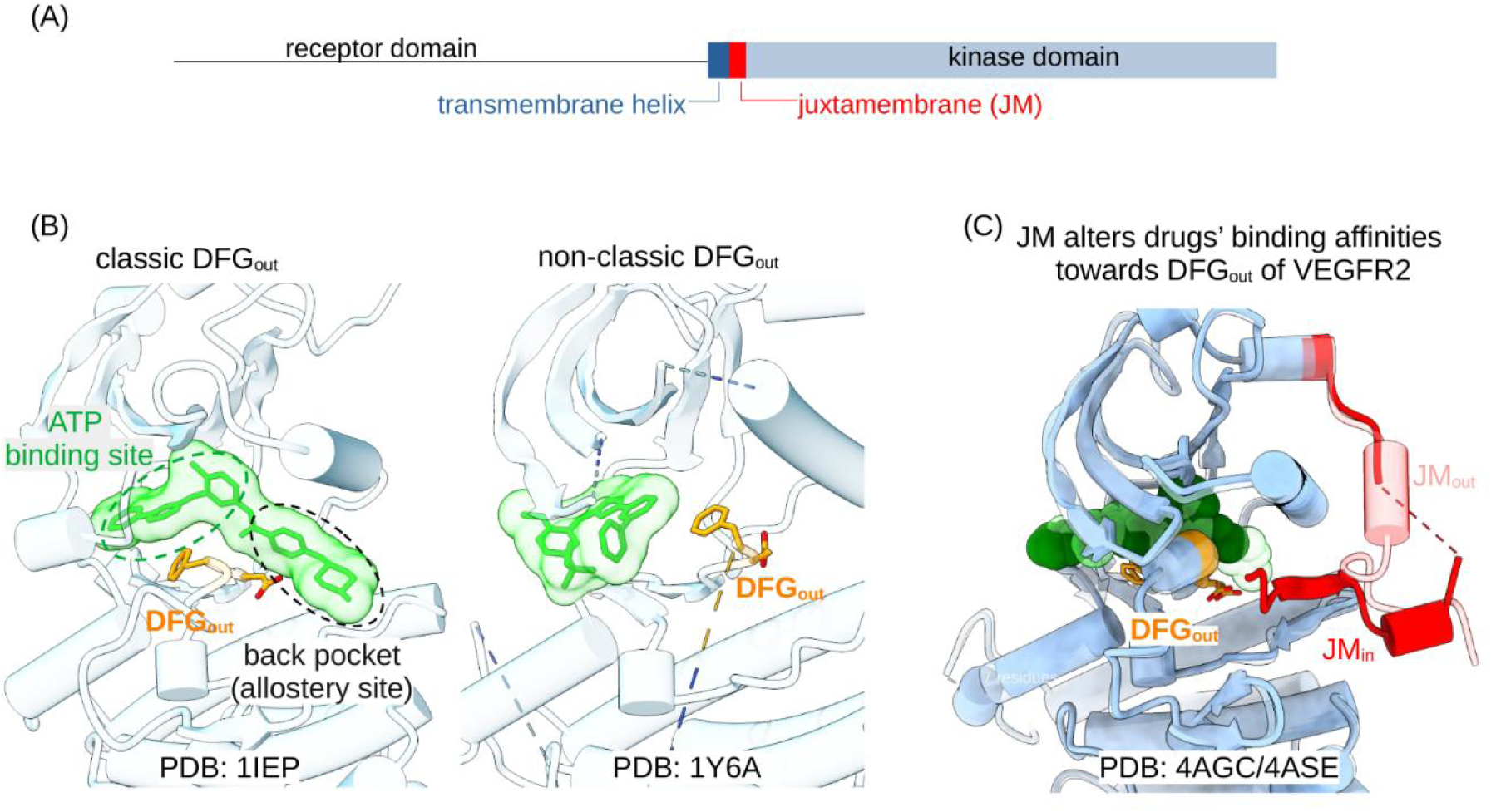
Structural background. (A) Domain organization of RTKs. (B) A classical DFG*_out_*conformation (PDB 1IEP^[21]^) with a fully formed back pocket, compared to a non-classical DFG*_out_* conformation (PDB 1Y6A^[22]^) where the back pocket is absent or only partially formed. Inhibitors bound to these structures are shown as green sticks and surfaces. (C) The JM motif of VEGFR2 was reported to regulate type II inhibitor binding affinity toward the DFG*_out_*conformation through steric clashes, adopting JM*_in_* and JM*_out_*models^[20]^. In this work, using MD simulations, we show that the disordered JM regions in VEGFR2 and other RTKs play a conserved role in shifting non-classical DFG*_out_* conformations toward classical ones via transient contacts with the kinase domain. This effect is relevant to the JM-mediated regulation of inhibitor binding affinity toward VEGFR2.

The conserved DFG motif (Asp-Phe-Gly triad), located in the activation loop of kinases, plays a vital role in regulating kinase activity states and inhibitor binding modes. Depending on the aspartate residue’s orientation as pointing into the ATP site (DFG*_in_*) or away from it (DFG*_out_*), kinases adopt active or inactive conformations, respectively^[5]^. Inhibitors binding to the DFG*_in_* conformation are classified as type I, while those binding to the DFG*_out_* conformation are type II inhibitors. The DFG*_out_* conformation creates an extra back pocket adjacent to the ATP site for inhibitor binding (**Fig. 1B**). This back pocket contains kinase-specific features, making type II inhibitors generally more selective than type I inhibitors^[4]^. Therefore, targeting the DFG*_out_* conformation is a mainstream strategy for improving kinase selectivity in inhibitor design ^[6–9]^.

However, druggable DFG*_out_* structural resources are much less abundant than those for the DFG*_in_* conformation, as deposited kinase structures in the Protein Data Bank (PDB) are more enriched in DFG*_in_* conformations^[10]^. Moreover, the DFG*_out_* conformation is highly flexible; structural elements forming the binding site, such as the *α*C-helix, P-loop, and A-loop, can adopt diverse conformations in the inactive state^[10,11]^. Consequently, many variant DFG*_out_* conformations exist, and not all are structurally validated to accommodate a type II inhibitor^[12]^. Those DFG*_out_* conformations that can fully accommodate a type II inhibitor are referred to as the “classical DFG*_out_*” conformation by Vijayan et al^[12]^, characterized by a fully formed back pocket (**Fig. 1B**). The scarcity of DFG*_out_* structural resources, especially classical DFG*_out_* conformations with a formed back pocket, is a major obstacle for DFG*_out_*-targeted selective inhibitor design^[13,14]^. While factors affecting the kinase DFG*_in_*/DFG*_out_* flip such as the influences from gatekeeper residues^[5]^, the *α*C-helix^[15]^, and the protonation state of titratable residues^[16]^ have been revealed, the basis underpinning the plasticity of DFG*_out_* conformations and the occurrence of non-classical versus classical DFG*_out_* states is less explored. Gaining such knowledge could help expand the repertoire of druggable classical DFG*_out_* conformations for the design of selective RTK inhibitors.

The juxtamembrane (JM) motif in RTKs has recently been identified as a factor that can fine-tune the binding affinities of type II inhibitors. Located between the kinase domain and the transmembrane helix (**Fig. 1A**), the JM consists of around 40 or more residues and is intrinsically disordered^[17–19]^. McTigue et al reported that the JM of vascular endothelial growth factor receptor 2 (VEGFR2), an RTK involved in angiogenesis, can differentially regulate the affinities of drugs that bind to the DFG*_out_* conformation. Specifically, biochemical assays revealed that the presence of the JM altered the affinities of certain drugs (e.g., axitinib, sunitinib, pazopanib) while having a negligible effect on others (e.g., linifanib and sorafenib)^[20]^. However, the structural basis for how the disordered JM exerts this differential regulatory effect remains elusive. Although structural insights based on a subset of inhibitor-VEGFR2 co-crystal structures suggest the disordered JM can interfere with drug binding by switching between a JM*_in_* and a JM*_out_* configuration (**Fig. 1C**), such structural observations can hardly be extrapolated to other drugs. More importantly, given that the differential effects are exerted by the JM without inducing a flip from the DFG*_out_* to the DFG*_in_* conformation, it is tempting to hypothesize that the JM motif of VEGFR2 refines the sub-conformational space of the DFG*_out_* state, leveraging the intrinsic plasticity of DFG*_out_* conformations.

In this study, we performed extensive MD simulations to explore the impact of the disordered JM motif on the conformational space of the DFG motif in RTKs and its relationship to inhibitor binding at the active site of VEGFR2. We revealed that in VEGFR2, the disordered JM is highly dynamic and forms transient contacts with the kinase domain. Interestingly, these dynamic interactions exert a unique fine-tuning effect on the DFG*_out_* sub-conformational space, shifting populations from non-classical to classical DFG*_out_* conformations. This dynamic model provides the structural basis underpinning the reported regulatory effects of JM on inhibitor binding toward VEGFR2. More importantly, we demonstrated that in other RTKs beyond VEGFR2, the disordered JM similarly promotes classical DFG*_out_* conformations. Our data reveal that disordered JM domains in RTKs possess a conserved role in promoting classical DFG*_out_* conformation formation through dynamic interactions. This role is particularly favorable for creating druggable DFG*_out_* conformations that can be exploited for designing high-selectivity type II inhibitors.

## 2. Materials and Methods

### 2.1. Conventional MD simulations of VEGFR2 and other five RTKs

We performed conventional MD of the kinase domain of VEGFR2 with and without JM attached. The starting structures for both systems were PDB 4AGC^[20]^ in the DFG*_out_*conformation. The no JM system consists of residues 834 to 1168, while the with JM system having residues 790-1168. Missing regions in PDB 4AGC were modeled by copying and pasting AlphaFold^[23]^ predicted full length VEGFR2 structure (AF-P35968-F1). The structures were solvated in the center of a cubic box composed of TIP3P^[24]^ water molecules, with a minimum distance of 1.0 nm between the protein and the box boundaries. The salt concentration was set as 0.15 M NaCl, and the Amber ff99SB-ILDN^[25]^ force field was used. Energy minimization was carried out using the steepest descent algorithm with stopping criteria being either reaching maximum of 50,000 steps or the maximum force fell below 1000.0 kJ/mol/nm. The systems were subsequently equilibrated under the NVT ensemble for 100 ps to gradually raise and stabilize the temperature at 300 K. During this phase, positional restraints were applied to the heavy atoms of the protein with force constant of 1000 kJ/mol/nm^2^. Non-bonded interaction cutoff was set as 1.0 nm, and the long-range electrostatics were handled using the Particle Mesh Ewald (PME)^[26]^ method. Following NVT equilibration, the systems were further equilibrated under the NPT ensemble for 100 ps to adjust the density and stabilize the pressure at 1 bar. Pressure coupling was applied isotropically using the Parrinello-Rahman barostat^[27]^. The time step was set as 2 fs and bonds involving hydrogen atoms were constrained using the LINCS algorithm^[28]^. Starting from the equilibrated state, five independent 1 μs long production runs were initiated in the NPT ensemble at 300 K and 1 bar. The MD simulations were performed using GROMACS 2022^[29]^. We also performed 500 ns conventional MD on the isolated JM (residues 790-833) of VEGFR2 starting from full-length VEGFR2 structure (AF-P35968-F1) predicted by AlphaFold^[23]^.

For the other five RTKs (EPHA3, RET, ErbB4, PDFGRA and KIT), two conventional MD simulations were conducted for each of them, one considering JM and one without JM. The starting structures for these simulations are PDBs 4TWO^[30]^, 7DUA, 3BCE^[31]^, 8PQJ^[32]^, and 7ZW8^[33]^, respectively. The residue ranges of the with JM construct and no JM construct are listed in **Table S1**.

MD protocol are the same as the abovementioned protocol, and the production run length is 1 *μs*. MD simulations were performed using GROMACS 2022^[29]^, and all MD simulations conducted in this study were summarized in **Table S1**.

### 2.2. Well-tempered metadynamics

Based on the crystal structure 4AGC, we performed a 1 *μs* well-tempered metadynamics^[34]^ simulation using GROMACS 2022 coupled with PLUMED 2.8.2^[35]^ to explore the conformational landscape of the JM region attached to the kinase domain of VEGFR2. Two collective variables were defined: *R*1, the distance between the oxygen atom of Y801 and the nitrogen atom of L1049, and *R*2, the average distance between the oxygen and nitrogen atoms of V805 and the corresponding nitrogen and oxygen atoms of I1025. Biased potentials were deposited at a frequency of 0.5 ps with an initial Gaussian height of 1.2 kJ/mol and a bias factor of 10. Gaussian widths of 0. 1 nm were applied for both collective variables, which were restrained within a grid range of 0.0-12.0 nm for *R*1 and 0.0-10.0 nm for *R*2. All other simulation parameters, including the force field, water model, system setup, ion concentration, and protocols for energy minimization, heating, and equilibration, were consistent with those used in our previous conventional molecular dynamics simulations.

### 2.3. Free energy perturbation (FEP)

We performed FEP calculations to obtain the binding free energy ( ΔG) of seven small-molecule inhibitors (axitinib, sunitinib, tivozanib, sorafenib, cediranib, linifanib, and pazopanib) to the kinase domain of VEGFR2, both with and without the JM motif attached. For the construct without JM, the VEGFR2 construct comprised residues 806-1171, based on PDB 4AGC, while the JM-containing construct included residues 786-1171. For the JM-containing construct, we considered two scenarios: one with the JM adopting the static JM*_in_* configuration observed in PDB 4AGC, and another with a dynamic JM configuration sampled from MD simulations. Thus, for each inhibitor, three ΔG values were computed: one for the no-JM construct, one for the static JM*_in_*configuration, and one for the dynamic JM configuration. These calculated values were compared against experimentally reported values from^[20]^.

Prior to FEP calculations, the inhibitor-VEGFR2 complex structures were prepared (**Fig. S1**). Except for sunitinib, all complexes were obtained via molecular docking using AutoDock4^[36]^, with the top-scoring pose selected. For sunitinib, the docked pose was unstable during MD refinement, leading to solvent exposure and deviation from the binding pocket (**Fig. S2**). Therefore, the crystal structure binding mode from PDB 4AGD^[20]^ was used as the starting structure for FEP. Each inhibitor-protein complex was subjected to MD simulations to relax the structures. Small molecule parameters were derived from the GAFF^[37]^ force field, with point charges calculated using the AM1-BCC model^[38]^. The MD protocol followed the aforementioned approach, with TIP3P water and counterions (NaCl) added to neutralize the system. For the no-JM and static JM_in_ cases, 100 ns production MD simulations were performed per complex, while for the dynamic JM case, 500 ns production runs were conducted. Three snapshots were extracted from the MD trajectories to serve as starting points for FEP calculations (see **Table S1**).

The FEP decoupling process was regulated through three distinct λ-channels: restraint-λ, coupling-λ, and vdw-λ, which systematically eliminated different interaction components between the ligand and protein. The electrostatic parameter (coul-λ) transitioned rapidly in the early stage, with the electrostatic interaction gradually increased from being completely off to fully on from window 0 (λ = 0.0) to window 4 (λ = 1.00) following the sequence of 0.0 → 0.25 → 0.50 → 0.75 → 1.00, while λ was maintained at 1.00 from window 5 to 19 to stabilize the electrostatic environment; the Van der Waals (vdW) parameter (vdW-λ) underwent a gradual transition after the electrostatic environment was stabilized, with λ kept at 0.00 (no vdW interaction) from window 0 to 4, and the vdW interaction fully activated from window 5 (λ = 0.05) to 19 (λ = 1.00) through a gradient increment following the sequence of 0.05 → 0.10 →… → 0.95 → 1.00; the restraint parameter (restraint-λ) was gradually strengthened as the calculation progressed, with λ set to 0.0 (no restraint, allowing the system to adapt freely) from window 0 to 1, the restraint strength increased in a gradient manner from window 2 (λ = 0.10) to 10 (λ = 0.90), and λ fixed at 1.0 (maximum restraint) from window 11 to 19 to ensure conformational stability during the later vdW transition stage. A schematic illustration for these λ settings are provided in **Fig. S3**, and we inspected the overlap in λ-space during the FEP calculation to validate the choice of λ scheme. Each window was simulated for 2 ns with a 2 fs timestep. The binding free energy was calculated using the Bennett acceptance ratio (BAR) method^[39]^. Each reported ΔG value represents the average of three independent FEP calculations using different inhibitor-protein complex structures.

### 2.4. Analysis

Standard analyses, including RMSF, SASA and secondary structure, distance, and angle calculations, were performed using the CPPTRAJ program^[40]^. Difference contact network analysis (dCNA) was carried out using the package from Yao et al^[41]^. This method first calculates the residue-residue contact frequency from the MD trajectories of the VEGFR2 kinase domain without and with the JM motif. A contact was defined as any pair of non-hydrogen atoms within a distance cutoff of 4.5 Å. A consensus contact network was constructed by including only contacts with a formation probability exceeding 0.9 in both systems. The Girvan-Newman algorithm^[42]^ was applied to this consensus network to automatically partition the protein into communities (functional domains), ensuring a consistent basis for comparison. A residue-wise difference contact network was then built from the change in contact probability between the two systems. In this step, only non-local residue pairs (sequence separation >3 residues) were considered to focus on non-covalent, allosterically relevant interactions. Finally, these residue-level differences were mapped onto the predefined communities to generate a coarse-grained community-community difference network. This network quantifies the net changes in interactions between protein domains, enabling a functional interpretation of the conformational dynamic changes induced by the JM motif. The volume of the inhibitor binding pocket was measured using the MDpocket tool^[43]^. The measurement employed the program’s default parameters, including a grid spacing of 1.0 Å and a probe radius of 1.4 Å to simulate the vdW surface. Plotting was performed using Matplotlib, and structural rendering was done with PyMOL, VMD and ChimeraX. Scripts used for this project are available at: https://github.com/bsu233/bslab/tree/main/2025RTKs.

## 3. Results

### 3.1. Disordered JM motif promotes classical DFG_out_ conformation formation in VEGFR2 by forming transient contacts with the kinase domain

We first verified the intrinsically disordered nature of the JM motif in VEGFR2. In its isolated state, the JM region was predicted to be almost entirely disordered by PONDR^[44]^. This was confirmed by 500 ns of atomistic MD simulations, which revealed that only few residues formed *α*-helical secondary structure, while the majority adopted coil or bend conformations (**Fig. 2A**). When attached to the kinase domain, the JM motif maintained its disordered and dynamic nature, sampling multiple conformations on the kinase surface during 5 × 1 *µs* MD simulations (**Fig. 2B**). The time-dependent formation and breakage of hydrogen bonds between the JM and kinase domains demonstrated that their interactions are transient rather than stable (**Fig. 2B**).

**Figure 2.**
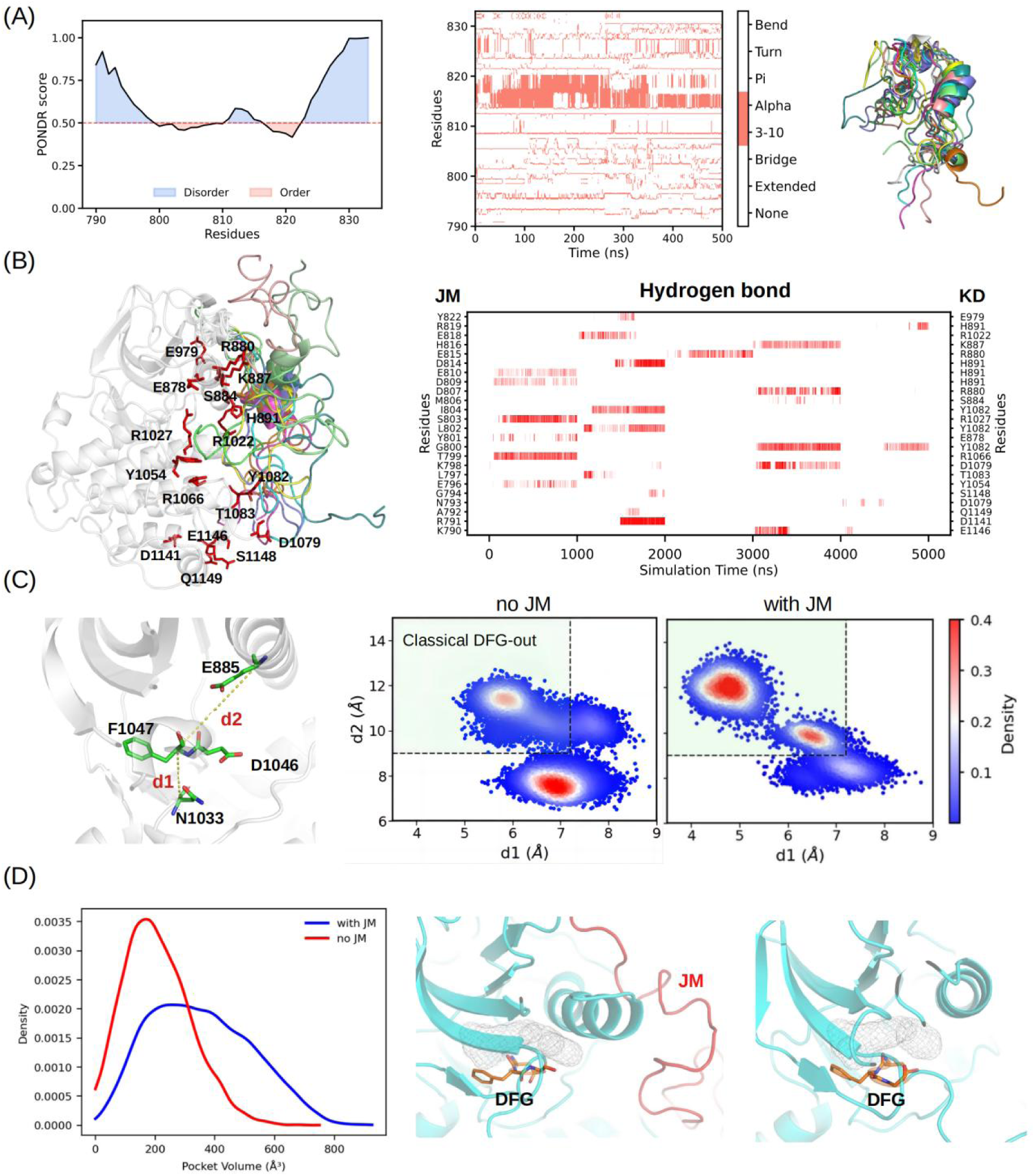
JM promotes classical DFG*_out_* conformation formation in VEGFR2. (A) PONDR-predicted disorder score, and the secondary structure of the JM motif from MD simulations, with representative conformations shown. (B) Time-dependent hydrogen bond formation between the JM and kinase domains over 5 × 1 *μs* MD simulations. A red line indicates the presence of a hydrogen bond. Key residues are shown as red sticks, and representative conformations of JM from structural clustering are depicted as colored cartoons. (C) Definition of the classical DFG*_out_* conformation using the *d*1 and *d*2 distances proposed by Vijayan et al^[12]^, and a projection of MD-sampled conformations onto the *d*1-*d*2 space. (D) Binding site volume measured from MD trajectories of the kinase domain with and without the JM motif attached.

To explore how these transient interactions affect the kinase domain, we performed 5 × 1 *µs* MD simulations of the VEGFR2 kinase domain in the absence of the JM motif. Both systems (with and no JM) were started from the DFG*_out_*conformation based on PDB 4AGC. We used the definition of a “ classical DFG*_out_*” conformation proposed by Vijayan et al^[12]^, which features a fully formed back pocket. This is quantified by two distances: *d*1, the Cα distance between the Asn in the HRDxxxxN motif and the Phe in the DFG motif, and *d*2, the Cα distance between the conserved Glu in the αC-helix and the DFG-Phe (**Fig. 2C**). A classical DFG*_out_* conformation is defined by d1 < 7.2 Å and d2 > 9.0 Å. Projecting our MD simulations onto the d1-d2 plane revealed that the DFG motif sampled a broad range of conformations, reflecting the intrinsic plasticity of the VEGFR2 DFG*_out_*state (**Fig. 2C**). Importantly, in the presence of the JM motif, the DFG sampling was concentrated within the classical DFG*_out_*region. Compared to the JM-less simulation, a clear population shift from non-classical to classical DFG*_out_* conformations was observed. Notably, across our cumulative 10 *µs* of simulation time, we did not observe a flip to the DFG*_in_* state, consistent with the relatively high energy barrier (∼10 kcal/mol ^[45]^) reported for the DFG*_in_*/DFG*_out_* transition in c-MET kinase. Measurements of the binding site volume (**Fig. 2D**) and solvent-accessible surface area (**Fig. S4**) both confirmed that the presence of the JM motif expanded the binding site compared to the no-JM case. This provides direct evidence that classical DFG*_out_* conformations are sampled more frequently when the JM motif is present. **These data demonstrate that the intrinsically disordered JM domain shapes the DFG*_out_* conformational landscape by promoting sampling of the classical DFG*_out_* conformation through transient contacts with the kinase domain.**

### 3.2. Allosteric signaling network underpins JM’s effect on DFG*_out_* sub-conformational space

We next sought to understand how the dynamic JM domain exerts its influence over the DFG*_out_*sub-conformational space through transient interactions. Tyrosine kinase domains are characterized by their plasticity and rich functional dynamics, which enable them to respond to diverse stimuli^[ 11]^. To investigate the JM’s effect on kinase domain dynamics, we plotted the RMSF values for the kinase domain with and without the JM motif (**Fig. 3A**). Comparison revealed that several regions, predominantly on the periphery of the kinase domain, exhibited altered flexibility in the presence of JM. These regions included not only those proximal to the JM anchor point (as expected) but also others distant from it (**Fig. 3A**). The altered dynamics in distant regions suggest that the JM motif has a clear allosteric regulation effect on the kinase domain. Interestingly, while peripheral regions showed significant dynamic changes, the core region surrounding the ATP-binding site experienced only subtle alterations. This pattern that large changes in the outer layer and subtle changes in the inner core may represent an allosteric signaling signature of kinase domains. This signature could allow kinases to maintain the conserved functional dynamics necessary for catalytic activity while still responding to regulatory stimuli.

**Figure 3.**
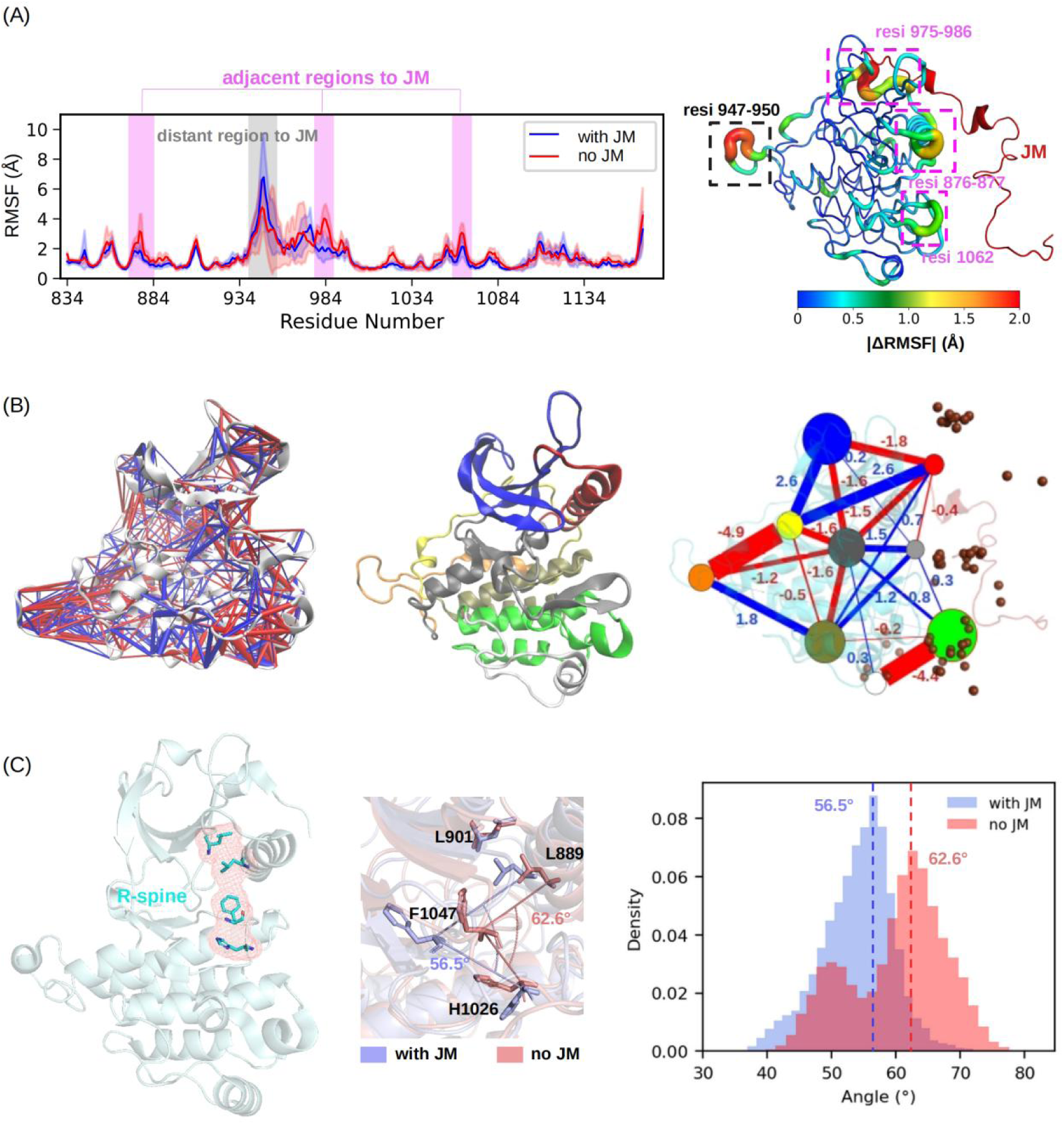
Allosteric signaling network within VEGFR2 kinase domain. (A) RMSF of kinase domain residues. The shaded area represents standard deviations calculated from five independent 1 *μs* MD trajectories. Residues with large RMSF differences are classified as adjacent or distant regions based on their spatial proximity to the JM’s position in the kinase structure. The absolute value of per-residue ΔRMSF (with JM minus no JM) is mapped onto the structure. (B) Residue contact difference network (with JM minus no JM) generated by the dCNA method^[41]^. The detailed contact map is shown in **Fig. S5**. Increased and reduced contacts are represented by blue and red lines, respectively, with line thickness proportional to the magnitude of change. The middle panel shows the nine defined groups/communities within the kinase domain used for community network analysis. The right panel quantifies the changes in signaling strength among these groups/communities using the dCNA method. The brown spheres depict the center of mass (COM) of JM sampled from MD simulations. (C) Effects of JM on the R-spine architecture of the VEGFR2 kinase domain. The straightness of the R-spine is quantified and compared between simulations with and without the JM motif.

To quantify the allosteric signaling changes induced by the JM motif, we employed difference contact network analysis (dCNA)^[41]^. This method constructs a network based on residue-residue contact changes between simulations with and without JM. Residues are then grouped according to protein domain organization to build a community-wide network, enabling the quantitative presentation of function-aware allosteric communication. As shown in **Fig. 3B**, the JM motif disrupts contacts among three key structural elements: the N-lobe *β*-sheet (blue node), the *α*C-helix (red node), and a core node bridging the N-lobe and C-lobe (dark gray node). The core node’s allosteric couplings with other nodes were more nuanced and complex than those among peripheral nodes. Specifically, the core node was connected by a greater number of links, and the associated magnitudes of these changes were smaller. This implies that the allosteric effect of the JM motif on the DFG*_out_* sub-conformational space is propagated through a layered signaling network. This intrinsic network allows the JM motif to fine-tune the DFG*_out_*sub-conformational space.

To demonstrate that the JM’s effect is limited to fine-tuning the DFG*_out_* sub-conformational space instead of inducing a DFG*_out_*-to-DFG*_in_* flip, we examined the structural state of the R-spine. The R-spine is a conserved structural motif in kinases, consisting of four hydrophobic residues, including the Phe from the DFG motif (**Fig. 3C**). The straightness of the R-spine is correlated with the DFG*_in_*/DFG*_out_*state and thus the activity of the kinase^[4]^. We show in **Fig. 3C** that the presence of the JM motif reduces the straightness of the R-spine. This is a signal that JM promotes the classical DFG*_out_* conformation rather than inducing a switch to the DFG*_in_* state. A similarly moderate effect of JM on another activity-related motif, the *α*C-helix^[46]^, was also observed (see **Fig. S4**). **The JM mottif allosterically regulates the kinase domain through a layered network. In this network, peripheral regions experience large-magnitude changes in coupling strength, while the core region exhibits more nuanced alterations. This network-mediated effect enables the fine-tuning of the DFG*_out_*sub-conformational space.**

### 3.3. Crystal structure-resolved JM*_in_* configuration in VEGFR2 is a ther-modynamically accessible metastable state of JM

We have demonstrated that the JM domain maintains a highly dynamic nature even when attached to the kinase domain. However, inhibitor-VEGFR2 co-crystal structures reported by McTigue et al suggest the JM can also bind into a cleft to form a stable, locked JM*_in_* conformation (PDB 4AGC and 4AGD^[20]^, see **Fig. S6**). To explore the relevance of this locked JM*_in_* configuration to the dynamic JM domain, we conducted metadynamics simulations to fully sample the conformational space of the JM. Our goal was to determine if the JM*_in_*state is thermodynamically accessible. In the crystal structure, the JM*_in_*configuration is stabilized by an antiparallel *β*-sheet between the JM and the kinase domain, specifically through the residue pairs V805-I1025 and Y801-L1049 (**Fig. 4A**). We therefore defined two reaction coordinates: *R*1, the distance between the oxygen atom of Y801 and the nitrogen atom of L1049, and *R*2, the average distance between the oxygen and nitrogen atoms of V805 and the corresponding nitrogen and oxygen atoms of I1025. We performed 1 *μs* of metadynamics simulation to compute the free energy landscape of the JM domain (**Fig. 4B**). The landscape revealed that the JM samples a broad range of low-energy conformations, with at least four distinct metastable states identified. Both kinase domain-contacting JM conformations (states **a, b**, and **c**) and a freely solvated JM state (state **d**) were found to be stable. The most stable basin (state **c**) itself contains multiple JM configurations interacting with the kinase domain. These data underscore the highly dynamic nature of the JM domain, which allows it to sample multiple metastable states. More importantly, the JM*_in_*configuration from the crystal structure was also captured in our simulation as state **a** (RMSD of JM is ∼3 Å). The free energy difference between this JM*_in_*-like state and the global minimum (state **c**) is approximately 8.0 kcal/mol. This confirms that while not the most stable state, the locked JM*_in_* conformation is indeed thermodynamically accessible to the dynamic JM domain. **Using metadynamics, we demonstrated that the JM*_in_* conformation is one of several metastable states accessible to the dynamic JM domain.**

**Figure 4.**
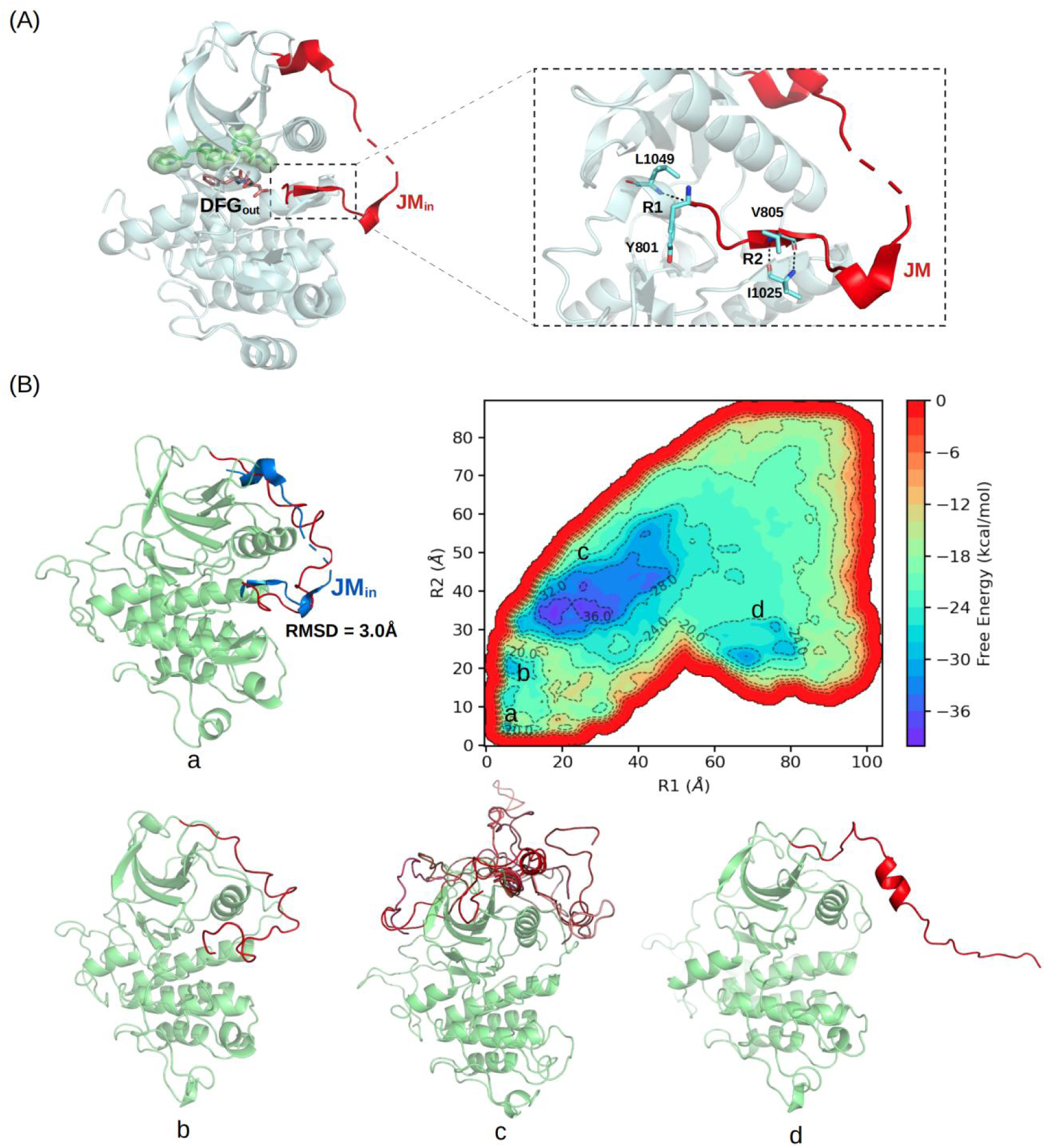
Conformational space of JM. (A) Reported JM*_in_* conformation in PDB 4AGC^[20]^ in which the JM docks back to the cleft and forms hydrogen bonds with L1049 and I1025 residues. Two reaction coordinates *R*1 and *R*2 were defined for metadynamics simulations. (B) The free energy landscape of JM conformational space. Four metastable states (**a, b, c**, and **d**) were identified. The overlap of state **a** with the reported JM*_in_* conformation (PDB 4AGC) was shown.

### 3.4. Incorporating the dynamic JM model completes the structural explanation for JM’s regulatory effect on VEGFR2 inhibitor binding

McTigue et al^[20]^ experimentally demonstrated that the JM motif of VEGFR2 differentially regulates the binding affinities of several inhibitors toward the DFG*_out_*conformation. Briefly, JM enhanced the binding affinities for axitinib, sunitinib, pazopanib, cediranib, and tivozanib relative to the no-JM construct^[20]^, while having negligible effects on linifanib and sorafenib. In co-crystal structures, JM adopted the JM*_in_* configuration with axitinib and the JM*_out_* configuration with tivozanib (**Fig. S6**), suggesting a correlation between JM conformational states and drug binding. Since we previously showed that the dynamic JM represents the major stable state while JM*_in_* is metastable, we sought to investigate how these JM states correlate with inhibitor binding affinities.

We conducted FEP calculations to determine inhibitor binding free energies (ΔG) to VEGFR2 using three constructs: no-JM, static JM*_in_*, and dynamic JM (**Fig. 5A**). Validation against experimental values for the no-JM construct showed good agreement for all inhibitors except sunitinib (**Fig. 5B**). The discrepancy for sunitinib likely stems from its hydrophobic tail, which becomes substantially solvent-exposed and unstable in the VEGFR2 binding site (**Fig. S2**), exhibiting large structural fluctuations during MD refinement that challenge FEP accuracy. After excluding sunitinib, FEP-calculated ΔG values showed an RMSE of 1.76 kcal/mol against experimental measurements, validating our approach. Using a static JM*_in_* model, the RMSE increased to 2.43 kcal/mol, indicating substantial discrepancy. The dynamic JM model showed slightly better correlation with an RMSE of 2.12 kcal/mol. Both JM-containing models exhibited RMSE >2 kcal/mol, suggesting neither alone accurately captures experimental affinities. Notably, when we selected for each inhibitor the closest-to-experimental ΔG value from either the JM*_in_* or dynamic JM models, the calculated values showed excellent agreement with experiments (RMSE = 1.42 kcal/mol; see FEP(mix JM) data in **Fig. 5C**). **These findings indicate that JM conformational states correlate with inhibitor binding affinities, and incorporating both dynamic and static JM*_in_* models provides a more complete explanation of JM’s regulatory effects on drug binding to the DFG*_out_*conformation of VEGFR2.**

**Figure 5.**
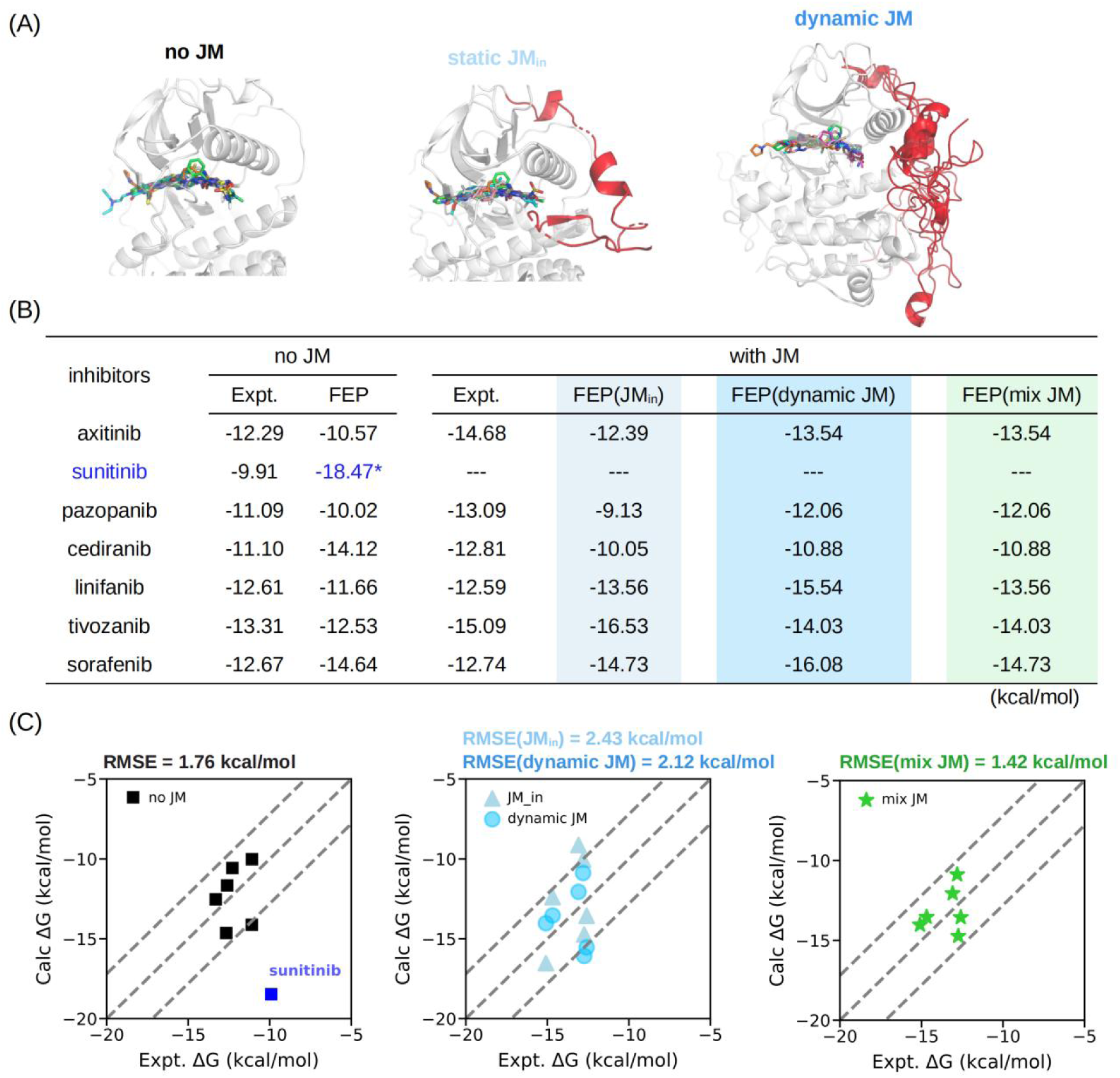
FEP-calculated inhibitor. Δ**G with VEGFR2.** (A) Three VEGFR2 constructs were considered: the no-JM construct, the reported JM*_in_* construct (PDB 4AGC), and a dynamic JM construct. (B) Tabulation of FEP-calculated ΔG and experimental ΔG values^[20]^. Each calculated ΔG represents the average of three independent runs. Sunitinib was excluded from the comparison as its FEP-calculated ΔG significantly deviated from the experimental value, likely due to its large solvent-exposed hydrophobic tail in the binding mode (see **Fig. S2** for details). The “ mix JM” column shows the calculated ΔG value (from either the JM*_in_* or the dynamic JM model) that is closest to the experimental result. (C) Correlation between FEP-calculated ΔG and experimental ΔG values. The dash line represents a deviation of ±2 kcal/mol.

### 3.5. Conserved role of JM in fine-tuning DFG*_out_* sub-conformational space to promote classical DFG*_out_* conformation formation across RTKs

The JM motif is highly conserved across RTKs. To investigate whether JM motifs in other RTKs similarly influence the DFG*_out_* sub-conformational space, we selected five RTKs whose JM motifs have been structurally resolved. The sequences of these JM motifs are highly conserved, and their PDB structures are shown in **Fig. 6A-B**. Starting from these structures, we performed 1 *μs* atomistic MD simulations for each RTK, both with and without the JM motif attached to the kinase domain. Notably, among the selected RTKs, three (EPHA3, RET, and ErbB4) have a DFG*_in_* configuration in their crystal structures, while the other two (PDGFRA and KIT) have a DFG*_out_* configuration. As shown in **Fig. 6C-D**, the JM motif affected the DFG conformational space in all five RTKs. The projections of the DFG motif onto the *d*1-*d*2 plane^[12]^ differed between simulations with and without the JM motif. Interestingly, for systems starting from the DFG*_in_* configuration, while JM affected DFG sampling, this effect did not induce a DFG*_in_*-to-DFG*_out_* flip. Meanwhile, for the two RTKs that started in the DFG*_out_* configuration, the presence of the JM motif induced a population shift towards the classical DFG*_out_* conformation, similar to the effect observed in VEGFR2. **We demonstrate that the JM motif in RTKs fine-tunes the DFG conformational space, with its nuanced effect on the DFG*_out_* sub-conformational space shifting populations from non-classical to classical DFG*_out_* conformations.**

**Figure 6.**
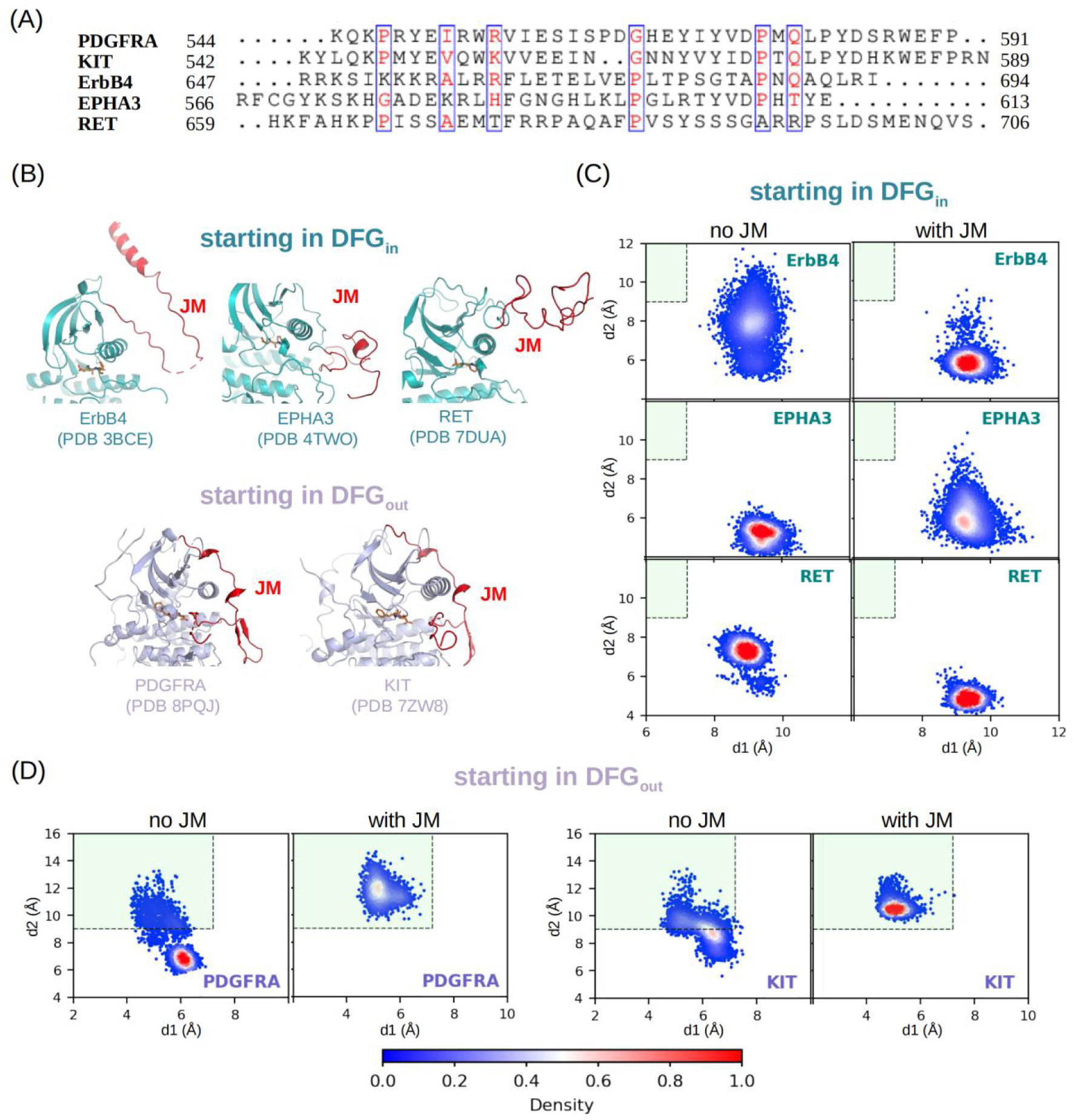
Effects of JM on the DFG conformational space in other five RTKs. (A) Sequence conservation of JM across five RTKs including PDGFRA, KIT, ErbB4, EPHA3, and RET. (B) The resolved crystal structure of these five RTKs are either in the DFG*_in_* conformation (ErbB4, EPHA3 and RET) or DFG*_out_* conformation (PDGFRA and KIT). For each RTK, 1 *μs* atomistic MD simulation was conducted for the kinase domain with and without JM attached, respectively. (C-D) MD samplings projection onto the *d*1-*d*2 plane to illustrate the effects of JM on DFG conformational space. The light green area corresponds to classical DFG*_out_* conformations.

## 4. Discussion

Our work exploring the impact of the JM motif on DFG conformational space was partially motivated by the experimental work of McTigue et al^[ 20]^. They reported that the presence of the JM motif can differentially alter inhibitors’ binding affinities toward the DFG*_out_* conformation of VEGFR2 without inducing a DFG*_out_*-to-DFG*_in_* flip. This effect was originally ascribed to a steric JM*_in_* model, in which JM adopts a specific configuration that creates differential steric clashes with various drugs, thereby interfering with drug binding. However, through MD simulations, we demonstrated that the JM is intrinsically disordered and primarily forms transient contacts with the kinase domain. This finding challenges the explanatory power of a purely static model for JM’s regulatory effect on inhibitor binding. Surprisingly, we discovered that transient interactions between the JM and kinase domain fine-tune the DFG*_out_* conformation of VEGFR2, particularly by promoting the formation of classical DFG*_out_* conformations. This effect enlarges the binding site relative to the no-JM construct (**Fig. 2**). Using FEP calculations, we demonstrated that incorporating the dynamic JM model allows accurate reproduction of experimental binding affinities of the JM-containing VEGFR2 constructs, achieving an RMSE of 1.42 kcal/mol. These results indicate that both dynamic JM and JM*_in_*models contribute to regulating inhibitor binding relative to the no-JM construct, with model preference being case-dependent.

Although we have revealed that the JM motif can affect the DFG conformational space, we emphasize that this effect is more prominent within the DFG*_out_*sub-conformational space than in altering the overall DFG*_in_* and DFG*_out_* conformation distributions. Specifically, we demonstrated that JM’s effect on the DFG conformational space in RTKs depends on the initial DFG state. When the initial state is DFG*_in_*, JM does not induce a DFG*_in_*-to-DFG*_out_* flip. Similarly, in simulations starting from DFG*_out_*, we observed no JM-induced DFG*_out_*-to-DFG*_in_* transition. These observations are consistent with the reported energy barrier of ∼ 10 kcal/ mol separating DFG*_in_* and DFG*_out_*conformations in some kinases^[45]^. Within the DFG*_out_* framework, however, the dynamic JM prominently shifts the sub-conformational space toward classical DFG*_out_* conformations. The existence of multiple DFG*_out_* conformations implies that the energy landscape underlying the DFG*_out_* sub-conformational space is rugged, making it susceptible to JM’s dynamic influence. We propose that the thermodynamic effect exerted by JM is smaller in magnitude than the energy barrier separating DFG*_in_* and DFG*_out_* conformations, but comparable to those separating sub-metastable states within the DFG*_out_* conformational space. Nevertheless, the nuanced classical DFG*_out_* conformation-promoting effect of JM revealed here is relevant for targeted type II inhibitor design against RTKs, as both in silico modeling of druggable DFG*_out_* conformations and experimental inhibitor profiling would benefit from using JM-containing constructs.

The mutual selection between inhibitor type and DFG conformation is complex. Multiple factors including differences among kinase subfamilies, local mutations within the DFG cleft, and distant residues in the kinase domain contribute to the preference of inhibitors for either the DFG*_in_*or DFG*_out_* conformation. The VEGFR2 inhibitors studied in this work all bind to the DFG*_out_* conformation^[20]^, yet they can also bind potently to the DFG*_in_* conformations of other non-RTK enzymes. For example, axitinib has been repurposed as a potent inhibitor for a drug-resistant mutant of the BCR-ABL1 kinase^[47]^, binding to the DFG*_in_* conformation of wild-type BCR-ABL1 and to the DFG*_out_*conformation of the BCR-ABL1 mutant. A similar phenomenon has been observed for sunitinib, which binds to the DFG*_in_* conformation of PAK6 kinase (a non-RTK)^[48]^. Additionally, numerous other factors have been reported to influence kinase selectivity. For instance, distant regulatory residues and binding pocket reorganization contribute to inhibitor selectivity across kinases^[49,50]^. Moreover, ligand binding can induce specific interactions outside the binding cleft that reciprocally modulate binding affinity^[51]^. Given this context, although we have revealed a conserved role of the JM motif in promoting classical DFG*_out_* conformations in RTKs, translating JM’s effect into general selectivity determinants for inhibitors requires caution.

## 5. Conclusions

In the present study, we performed MD simulations and FEP calculations to explore the impact of the disordered JM motif on the conformational space of the DFG motif in VEGFR2 and other RTKs, and to relate this impact to inhibitor binding to VEGFR2. We demonstrated that the JM motif is highly dynamic and forms transient interactions with the kinase domain of VEGFR2. Despite their transient nature, these interactions can shape the DFG*_out_* conformational space of VEGFR2, shifting populations from non-classical to classical DFG*_out_* conformations. This ability of the JM motif is realized through a layered allosteric signaling network within the kinase domain of VEGFR2. This layered network transmits the substantial dynamic changes occurring at the periphery of the kinase domain (that induced by JM interactions) into nuanced and sophisticated modulations of the core region, ultimately fine-tuning DFG*_out_* conformations. FEP calculations further demonstrate that this dynamic JM model is relevant to the experiment measured inhibitors affinities toward the JM-containing construct of VEGFR2. Finally, we showed that in other RTKs, the conserved JM motif similarly promotes the formation of classical DFG*_out_* conformations, as observed in VEGFR2. Overall, our findings suggest that using JM-containing RTK constructs is beneficial for modeling druggable DFG*_out_*conformations and for profiling the biochemical properties of type II inhibitors against RTKs.

## 6. Acknowledgements

Research reported in this work was supported by the Natural Science Foundation of China (32501103), Natural Science Foundation of Heilongjiang Province (PL2024B022), China Postdoctoral Science Foundation (2023M730827).

## Supplementary Information (SI)

**Figure S1.**
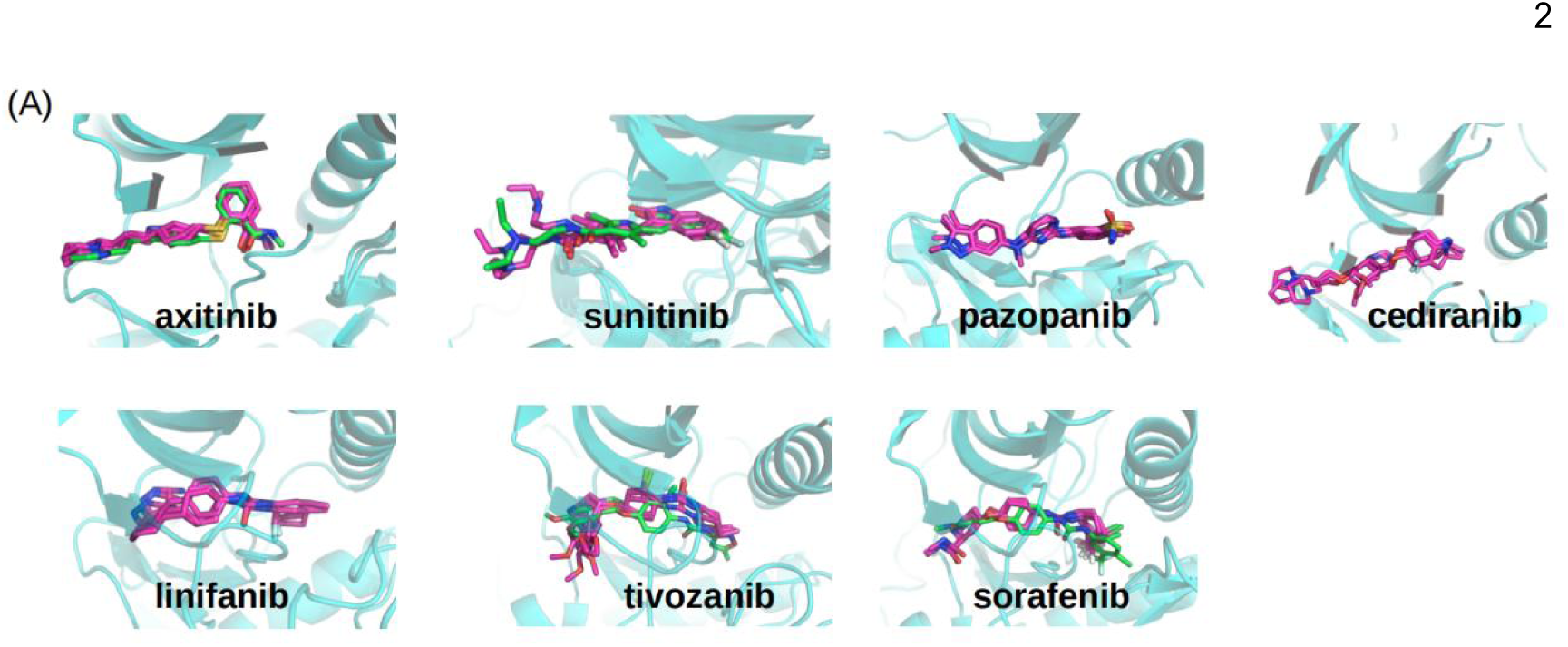
(A) The initial structures for FEP calculations were obtained through Autodock4^[ 36]^ docking for all complexes except sunitinib. The magenta stick representations in the figure show the ligand binding modes extracted from 93 ns, 96 ns, and 99 ns time points after 100 ns of molecular dynamics simulation of the docked complexes. The green stick representations correspond to the ligand binding modes from the crystal structures 4AGC^[20]^ (axitinib), 4AGD^[20]^ (sunitinib), 4ASE^[20]^ (tivozanib), and 4ASD^[20]^ (sorafenib).

**Figure S2.**
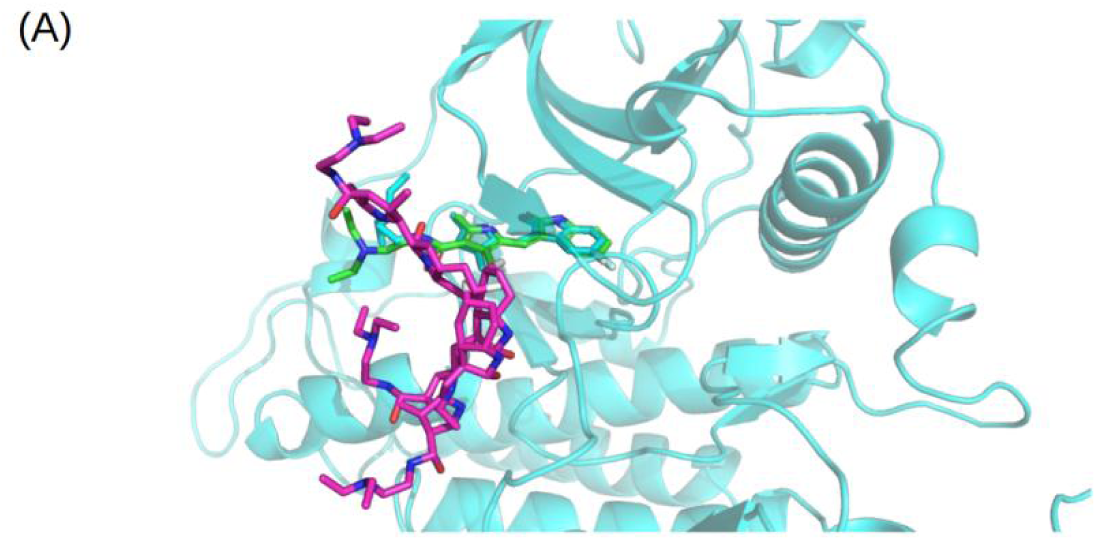
Analysis of the binding mode of Sunitinib. (A) The green stick model represents the binding conformation in the crystal structure, the cyan stick model represents the predicted binding mode from molecular docking, and the magenta stick model displays the conformations extracted at 93 ns, 96 ns, and 99 ns time points from the molecular dynamics (MD) simulation of the docked complex.

**Figure S3.**
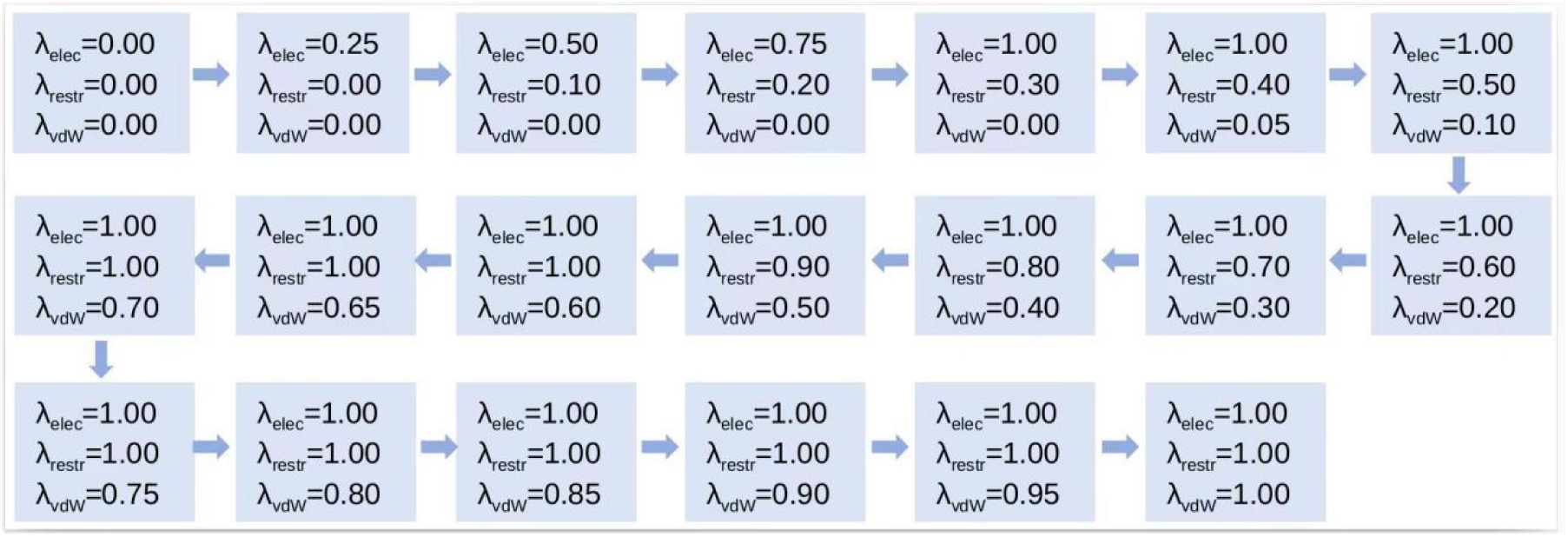
Schematic of λ setting in the FEP calculations.

**Figure S4.**
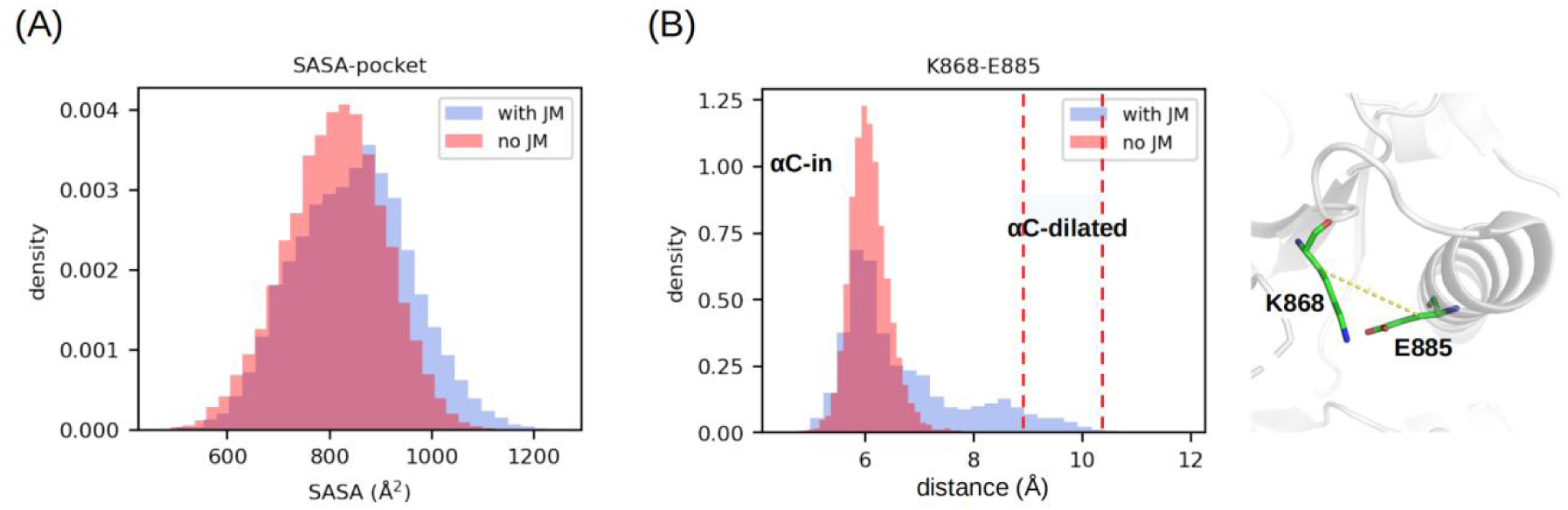
(A) Effects of JM on the solvent-accessible surface area (SASA) of the binding pocket, and (B) on *α*C-helix of VEGFR2. The *α*C-helix that has been reported as a key function regulation motif of kinase. We showed that the presence of JM redistributes the *α*C-helix configurations from dominant canonical “in” state to span more broad configurations, with some *α*C-helix sample the *α*C- dilated configuration, an intermediate conformation between “ in” and “ out” states^[^ ^46]^. The inward and outward movement of the *α*C helix, which are common to protein kinases, are critical features of the active and inactive forms. In the active form, the *α*C helix is located at the “in” position^[51]^.

**Figure S5.**
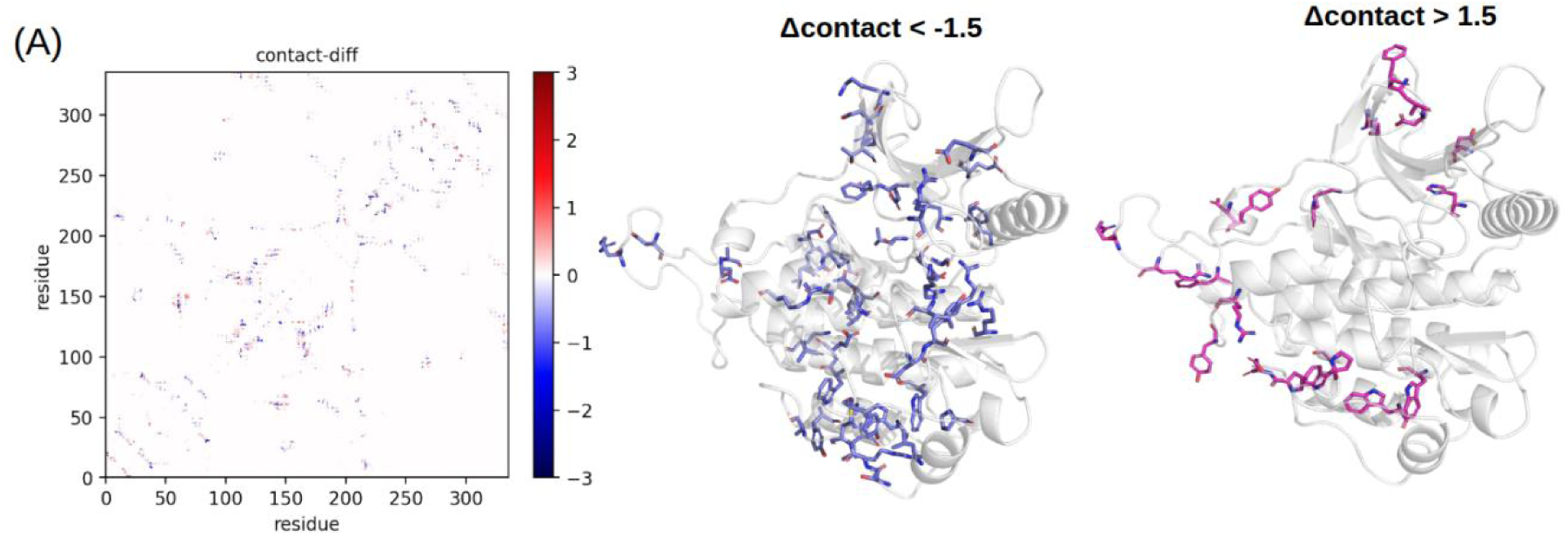
Contact map difference of the kinase domain of VEGFR2 (with JM minus without JM). Residue pairs that have significant Δcontact values are shown as sticks.

**Figure S6.**
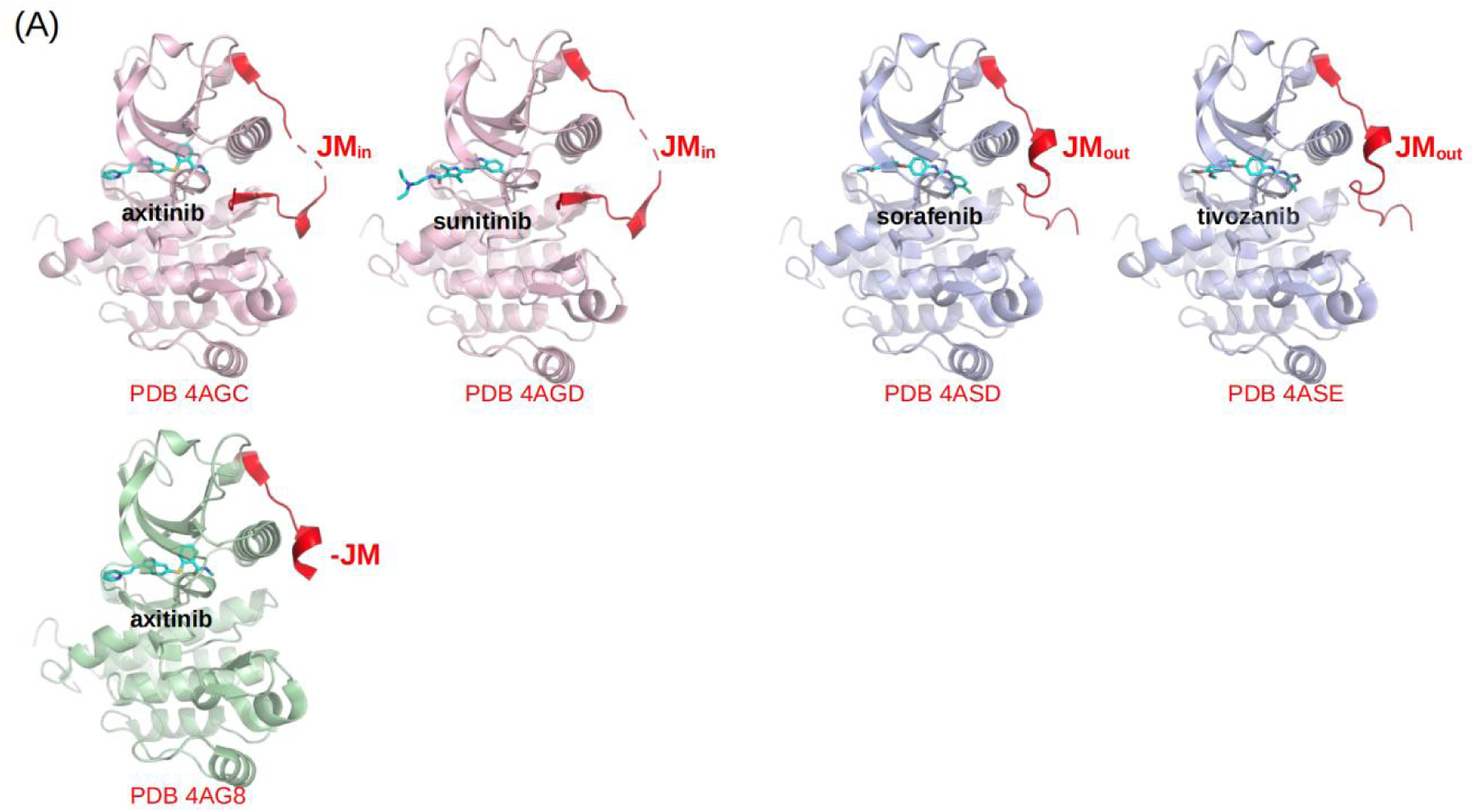
Reported inhibitor-VEGFR2 co-crystal structures by McTigue et al ^[20]^.

---

**Table S1.** Molecular dynamics simulations performed in this study.

| Simulation Types | Cases | Starting Structures | Simulation Length |
| --- | --- | --- | --- |
| **Conventional**<br>**All-Atom MD** | VEGFR2-JM | AF-P35968 (residues 790-833) | 500 ns |
| | VEGFR2 KD (with JM) | PDB 4AGC (residues 790-1168) | 1 μs × 5 |
| | VEGFR2 KD (no JM) | PDB 4AGC (residues 834-1168) | 1 μs × 5 |
| | EPHA3 KD (with JM) | PDB 4TWO (residues 566-906) | 1 μs |
| | EPHA3 KD (no JM) | PDB 4TWO (residues 604-906) | 1 μs |
| | RET KD (with JM) | PDB 7DUA (residues 661-1011) | 1 μs |
| | RET KD (no JM) | PDB 7DUA (residues 706-1011) | 1 μs |
| | ErbB4 KD (with JM) | PDB 3BCE (residues 651-966) | 1 μs |
| | ErbB4 KD (no JM) | PDB 3BCE (residues 688-966) | 1 μs |
| | PDGfra KD (with JM) | PDB 8PQJ (residues 550-973) | 1 μs |
| | PDGfra KD (no JM) | PDB 8PQJ (residues 590-973) | 1 μs |
| | KIT KD (with JM) | PDB 7ZW8 (residues 546-932) | 1 μs |
| | KIT KD (no JM) | PDB 7ZW8 (residues 588-932) | 1 μs |
|  | VEGFR2-inhibitors<sup>a</sup> (with JM) | PDB 4AGC (residues 786-1171) | 100 ns<sup>b</sup> |
|  | VEGFR2-inhibitors<sup>a</sup> (no JM) | PDB 4AGC (residues 806-1171) | 100 ns<sup>b</sup> |
|  | VEGFR2-inhibitors<sup>a</sup> (dynamic JM) | PDB 4AGC (residues 786-1171) | 500 ns<sup>c</sup> |
|  | inhibitors<sup>a</sup> | — | 100 ns<sup>d</sup> |
| **MetaDynamics** | VEGFR2 (with JM) | PDB 4AGC (residues 801-1168) | 1 μs |
| **Free Energy**<br>**Perturbation** | VEGFR2-inhibitors<sup>a</sup> (with JM) | PDB 4AGC (residues 786-1171) | 2 ns/$\lambda$ × 20 $\lambda$ × 3<sup>b</sup> |
| | VEGFR2-inhibitors<sup>a</sup> (no JM) | PDB 4AGC (residues 801-1171) | 2 ns/$\lambda$ × 20 $\lambda$ × 3<sup>b</sup> |
| | VEGFR2-inhibitors<sup>a</sup> (dynamic JM) | PDB 4AGC (residues 786-1171) | 2 ns/$\lambda$ × 20 $\lambda$ × 3<sup>c</sup> |
| | inhibitors<sup>a</sup> | — | 2 ns/$\lambda$ × 20 $\lambda$ × 1<sup>d</sup> |

<sup>a</sup> : axitinib, sunitinib, pazopanib, cediranib, linifanib, tivozanib, sorafenib.
<sup>b</sup> : Take the frames at 93, 96, and 99 ns from the CMD simulation as starting structures for FEP.
<sup>c</sup> : The 500 ns MD trajectory was subjected to cluster analysis, and the central conformations of the three most populated clusters were selected as the starting structures for FEP simulations.
<sup>d</sup> : Take the last frame of the simulation as the starting structure for FEP calculation.

